# Genomic surveillance of carbapenem-resistant *Klebsiella* in Wales reveals persistent spread of *K. pneumoniae* ST307 and adaptive evolution of pOXA-48-like plasmids

**DOI:** 10.1101/2022.05.25.492139

**Authors:** Sophia David, Massimo Mentasti, Kirsty Sands, Edward Portal, Lee Graham, Joanne Watkins, Catie Williams, Brendan Healy, Owen B. Spiller, David M. Aanensen, Mandy Wootton, Lim Jones

## Abstract

Rising rates of multi-drug resistant *Klebsiella* infections necessitate a comprehensive understanding of the major strains and plasmids driving spread of resistance elements. Here we analysed 540 *Klebsiella* isolates recovered from patients across Wales between 2007 and 2020 using combined short- and long-read sequencing approaches. We identified resistant clones that have spread within and between hospitals including the high-risk strain, sequence type (ST) 307, which acquired the *bla*_OXA-244_ carbapenemase gene on a pOXA-48-like plasmid. We found evidence that this strain, which caused an acute outbreak largely centred on a single hospital in 2019, had been circulating undetected across South Wales for several years prior to the outbreak. In addition to clonal transmission, our analyses revealed evidence for substantial plasmid spread, mostly notably involving *bla*_KPC-2_ and *bla*_OXA-48-like_ (including *bla*_OXA-244_) carbapenemase genes that were found among many species and strain backgrounds. Two thirds (20/30) of the *bla*_KPC-2_ genes were carried on the Tn*4401a* transposon and associated with IncF plasmids. These were mostly recovered from patients in North Wales, reflecting an outward expansion of the plasmid-driven outbreak of *bla*_KPC-2_-producing *Enterobacteriaceae* in North-West England. 92.1% (105/114) of isolates with a *bla*_OXA-48-like_ carbapenemase carried the gene on a pOXA-48-like plasmid. While this plasmid family is highly conserved, our analyses revealed novel accessory variation including integrations of additional resistance genes. We also identified multiple independent deletions involving the *tra* gene cluster among pOXA-48-like plasmids in the ST307 outbreak lineage. These resulted in loss of conjugative ability and signal adaptation of the plasmids to carriage by the host strain. Altogether, our study provides the first high resolution view of the diversity, transmission and evolutionary dynamics of major resistant clones and plasmids of *Klebsiella* in Wales and forms an important basis for ongoing surveillance efforts.

**Data Summary:** All raw short read sequence data and hybrid assemblies are available in the European Nucleotide Archive (ENA) under project accession PRJEB48990.

## Introduction

The *Klebsiella* genus, which belongs to the *Enterobacteriaceae* family, includes several species that are important opportunistic pathogens of humans. In particular, *Klebsiella pneumoniae* is a leading cause of healthcare-associated infections globally (Magill et al. 2014; Ling et al. 2015; Vincent et al. 2020). *Klebsiella* species cause a range of disease types including pneumonia, skin and wound infections, urinary tract infections, and sepsis. These occur most frequently in the elderly, neonates and immunocompromised individuals (Meatherall et al. 2009).

*Klebsiella* are commonly carried in the gastrointestinal tract of healthy humans (Podschun & Ullmann, 1998). Analysis using whole genome sequencing (WGS) performed on paired screening swabs and clinical samples has demonstrated that gastrointestinal carriage is a major reservoir of *K. pneumoniae* infections (Gorrie et al. 2017). In addition, outbreaks are frequently reported in healthcare settings, driven by transmission via person-to-person contact, contaminated environmental surfaces and medical devices (e.g. ventilators, catheters) (Snitkin et al. 2012; Gu et al. 2018; Tavoschi et al. 2020).

Across the world, studies have reported an increasing prevalence of infections with multidrug-resistant *Klebsiella* (Cassini et al. 2019; Castanheira et al. 2019). A particular concern has been the rise of infections that are resistant to carbapenems, which has been listed as a “critical” threat by the World Health Organisation (WHO, 2017). Carbapenems are a class of antibiotics that are vital for treatment of severe *Klebsiella* infections due to their broad spectrum of activity (e.g. against extended-spectrum beta lactamase (ESBL) producers) and limited adverse effects. Carbapenem resistance in *Klebsiella* is usually conferred by plasmid-encoded carbapenemase genes including *bla*_KPC_, *bla*_OXA-48-like_, *bla*_VIM_, *bla*_NDM_ and *bla*_IMP_ (Queenan & Bush, 2007). These are often associated with mobile genetic elements (MGEs) such as transposons, which commonly insert into plasmid backbones.

A new cohort of drugs has become available in recent years to treat carbapenem-resistant infections including novel beta lactam-beta lactamase inhibitor (BL-BLI) combinations (e.g. ceftazidime-avibactam, meropenem-vaborbactam) and the siderophore cephalosporin, cefiderocol. However, some of these drugs are not effective against certain types of carbapenemase producer, including the BL-BLIs which have no activity against metallo-betalactamases (*bla*_VIM_, *bla*_NDM_ and *bla*_IMP_). These also may not work reliably against non-carbapenemase-producing carbapenem resistant strains (Bradley & Lee, 2019).

As a result, a multi-pronged approach is required for the control of multi-drug resistant *Klebsiella*, including new therapeutics, heightened infection control and enhanced surveillance. The latter involves acquiring a detailed understanding of the diversity and transmission routes of different resistance mechanisms. Integration of genome sequencing into surveillance systems, which is ongoing across many public health laboratories, is providing a vital role in achieving this. However, while the current focus is largely on monitoring important strains and detecting clonal transmission, there is still a strong need for bioinformatic workflows that provide data on the role of specific plasmid vectors in resistance spread.

The Specialist Antimicrobial Chemotherapy Unit (SACU) based at Public Health Wales (PHW) Microbiology in Cardiff provides a reference laboratory service for the identification and confirmation of antimicrobial resistance mechanisms in bacterial isolates including *Klebsiella*, referred by microbiology laboratories across and beyond Wales. It also collects *Klebsiella (*and other bacterial) isolates from blood cultures for surveillance purposes and receives isolates from suspected outbreaks as part of the development of PHW’s Whole Genome Sequencing (WGS) based typing service, in partnership with the Pathogen Genomics Unit (PenGU). This study describes the retrospective genomic analysis of a collection of *Klebsiella* isolates (*n*=540) gathered by SACU between 2007 and 2020. Using a combination of short- and long-read sequencing, our analysis revealed detailed insights into the evolutionary dynamics of the major *K. pneumoniae* clones and plasmids driving spread of antibiotic resistance genes in Wales.

## Methods

### Sample collection

540 *Klebsiella* isolates were included in this study (**Supplementary Tables 1 & 2**). Isolates were obtained from clinical, screen or environmental samples between 2007 and 2020 and bead-stored at -80°C within the SACU culture collection at PHW. They were cultured overnight from beads on blood agar (OXOID, UK) at 35 ± 1°C prior to sequencing.

### Short read sequencing

Isolates were subbed onto 500μL of Tryptone Soya Broth (TSB) (E&O Laboratories, UK) and incubated overnight at 35 ± 1°C. Growth from 125μL of TSB culture was harvested at 12,000*g* for 2 min and the supernatant discarded. A lysis step was performed by adding 190μL of Buffer G2 (Qiagen, Germany) and 10μL of Proteinase K (Qiagen, Germany) followed by incubation at 56°C for 30 min. DNA was extracted from the lysates using a generic 3.0.4 protocol on the EMAG platform (bioMerieux, France) according to the manufacturer’s instructions and eluted in 100μL final volume. DNA was quantified using the FLUOstar Omega fluorometer (BMG Labtech, Germany) and normalised to approximately 0.4ng/μL. Sequencing libraries were prepared using the Nextera XT DNA library preparation kit and combinations of IDT index plates (Illumina, USA). The final libraries were cleaned through AMPure SPRI bead (Beckman Coulter, USA) fragment size selection, and quantified by the Collibri™ Library Quantification Kit (Invitrogen, USA) before normalising to approximately 0.5nM. The normalised final libraries were pooled and loaded onto MiSeq or NextSeq platforms (Illumina, USA).

### Long read sequencing

Isolates were subbed onto 2mL of LB broth (Becton Dickinson, UK) and incubated overnight at 35 ± 1°C. DNA was extracted using the automated QIAcube (Qiagen) platform with an additional RNAse step and then quantified using the Qubit 4.0 (Thermofisher) using HS and BR dsDNA kits as appropriate. DNA was purified and concentrated using SPRI beads (Mag-Bind TotalPure, Omega) at a 1:1 ratio with a final 15μL elution in nuclease-free water. Genomic libraries were prepared using the Rapid Barcoding Kit (SQK-RBK004; Oxford Nanopore Technology, UK). Eight to twelve isolates were loaded onto a FLO-MIN106 R9.4 flow cell and sequenced on a MinION device (Oxford Nanopore Technology, UK). Sequencing was performed on an Intel i7-6700 desktop computer over a 72 h period and basecalling was performed within MinKnow using Guppy v4.5.4 (Oxford Nanopore Technology, UK).

### Short-read and hybrid assembly

Short sequence reads from all isolates were assembled using SPAdes v3.10.0 (Bankevich et al. 2012) using the “--careful” flag and the “--cov-cutoff” flag set to “auto”. Long sequence reads were de-multiplexed within Minknow and assembled with the corresponding short reads using Unicycler v0.4.7 (Wick et al. 2017) with default parameters. Assembly statistics are available in **Supplementary Tables 1 and 2**. Assemblies were annotated using Prokka v1.14.5 (Seemann, 2014).

### Characterisation of *Klebsiella* genomes

Kleborate v2.0 (Lam et al. 2021) was used to determine the species and sequence type (ST) (for species with available schemes) from the assemblies, and identify virulence and resistance genes. Kaptive (Wyres et al. 2016), integrated into Kleborate, was used to type the capsule and O antigen biosynthesis loci.

### Species-wide phylogenetic analyses

Core genes (i.e. those present in ≥99% of isolates) were identified among assemblies of each *Klebsiella* species using Panaroo v1.2.4 (Tonkin-Hill et al. 2020) in the “sensitive” mode (in species with ≥5 isolates only). Variable positions were extracted from each core gene alignment using SNP-sites (Page et al. 2016). These were used to construct a maximum likelihood phylogenetic tree for each species using IQ-TREE v1.6.10 (Nguyen et al. 2015).

### Phylogenetic analyses of individual STs

Additional public genomes from *K. pneumoniae* ST307 (*n*=964) and ST1788 (*n*=2), the two most frequent STs in the PHW collection, were identified using the Pathogenwatch database (Argimon et al. 2021). Using the available accession numbers, all corresponding short-read data were downloaded from the ENA while the associated metadata was obtained from Pathogenwatch (https://pathogen.watch/).

Short-read sequencing data from ST307 and ST1788 isolates from both the PHW and public collections (including also single-locus variants of ST307 from the PHW collection) were mapped to ST-specific reference genomes. For ST307, this was the complete chromosome from the hybrid assembly of ARGID_32304, while the short-read assembly of ARGID_33748 (62 contigs) was used for ST1788. Mapping was performed using Burrows Wheeler Aligner (Li & Durbin, 2009) with SNPs identified using SAMtools v1.2 mpileup and BCFtools v1.2 (Li et al. 2009). Gubbins v2.4.1 (Croucher et al. 2015) was used to remove recombined regions from the resulting pseudo-genome alignments and generate maximum likelihood phylogenies based on the remaining variable sites. Pairwise SNP differences between isolates were determined from the pseudo-genome sequences using pairsnp (https://github.com/gtonkinhill/pairsnp).

### Visualisation of trees and metadata

Phylogenetic trees were visualised together with metadata using the Interactive Tree of Life tool v6 (Letunic & Bork, 2021) and further annotated using Adobe Illustrator v2017.1.0. Interactive visualisations were also created using Microreact v178 (Argimon et al. 2016).

### Characterisation of plasmids and MGEs

Plasmid replicon types carried by all isolates were determined from the short-read sequence data using Ariba v2.14.6 with the PlasmidFinder database (Carattoli et al. 2014). The PlasmidFinder tool (https://bitbucket.org/genomicepidemiology/plasmidfinder/src/master/) (Carattoli et al. 2014) was used to identify plasmid replicons in completely sequenced plasmids. oriTfinder v1.1 (Li et al. 2018) was used to identify the origin of transfer site (*oriT*) among complete pOXA-48-like plasmid sequences.

Linear comparisons between multiple pairs of complete plasmids were generated using EasyFig (Sullivan et al. 2011) and further annotated using Adobe Illustrator v2017.1.0. The percentage nucleotide similarities between individual plasmid genes and those from a reference plasmid were determined using BLASTn v2.6.0 (Camacho et al. 2009). The number of SNPs between pairs of aligned plasmids were determined using NUCmer v3.1 from the MUMmer package (Kurtz et al. 2004).

The presence of the Tn*4401* transposon among *bla*_KPC_-encoding isolates, together with the Tn*4401* variant type, was determined from the short sequence reads using TETyper v1.1 (Sheppard et al. 2018).

### Identification and comparison of pOXA-48-like plasmid sequences from short-read data

Sequence reads from all isolates were mapped to the 61.8kb IncL plasmid, pOXA48a (Poirel et al. 2012) (accession JN626286), using Burrows Wheeler Aligner (Li & Durbin, 2009) with SNPs identified using SAMtools v1.2 mpileup and BCFtools v1.2 (Li et al. 2009). The binary alignment map (BAM) file from each isolate was used to determine the length of the reference plasmid that was mapped by at least one sequence read. A SNP matrix, calculated from pseudo-plasmid sequences of isolates with ≥70% mapping to the reference, was used to construct a minimum spanning tree using the standalone version of GrapeTree v1.5.0 (Zhou et al. 2018). Pairwise SNP differences between all plasmid pseudo-sequences were also determined using pairsnp (https://github.com/gtonkinhill/pairsnp).

To determine the short-read plasmid coverage over the pOXA48a reference relative to the chromosomal coverage, we mapped all isolates to the NTUH-K2044 chromosome (accession AP006725.1) as described above. We used the “genomecov” function within bedtools v2.29.0 (Quinlan & Hall, 2010) to determine the chromosomal coverage using sorted BAM files as input, and determined the median value for each isolate. We then calculated the plasmid coverage, relative to the median chromosomal coverage, across 100bp windows of each mapped plasmid sequence (sliding every 20bp) and visualised the resulting matrix using the “heatmap.2” function in R v.4.0.2.

### Bacterial conjugation

Transfer of the pOXA-48-like plasmid-encoded *bla*_OXA-244_ gene by six *K. pneumoniae* ST307 isolates was determined using a liquid-mating conjugation method. All mating experiments were performed using both the *E. coli* J53 strain (200mg/L sodium azide) and a *K. pneumoniae* strain, 71.1 (passaged over 10d to generate a rifampicin MIC of 4,096mg/L) as separate recipients. Two mating culture ratios were tested, 1:3 and 5:1, for both donor and recipient respectively. Briefly, donor and recipient isolates were grown to log phase in LB broth and mixed according to the appropriate mating-culture ratio, then incubated at 37°C for 16-20h. Following serial dilution of the mating cultures, transconjugant colonies were selected for using appropriate chromogenic agar (Sigma Aldrich) and antibiotic combination (J53, 200mg/L sodium azide and 4mg/L meropenem; 71.1, 256mg/L rifampicin). To differentiate donor-recipient in the *K. pneumoniae-K. pneumoniae* experiment, the ST307 donor colonies were selected for using the chromogenic agar antibiotic combination (4mg/L meropenem and ampicillin at 32mg/L) as 71.1 was sensitive to ampicillin. Transfer of the *bla*_OXA-244_ was confirmed via PCR, and transconjugates from the *K*.*pneumoniae-K*.*pneumoniae* experiments were further confirmed by short-read (Illumina) sequencing. Briefly, gDNA from transconjugates was extracted using the automated QIAcube (Qiagen). gDNA libraries were prepared using the Nextera XT v2 kit with bead-based normalisation, and sequenced using V3 chemistry on an Illumina Miseq. The conjugation frequency was calculated as a ratio of the CFU/mL of transconjugate CFU/mL to the donor isolate.

### Data availability

All raw short read sequence data and hybrid assemblies are available in the ENA under project accession PRJEB48990. Individual accession numbers for samples are available in **Supplementary Tables 1 and 2**.

## Results

### High diversity of carbapenemase-producing *Klebsiella* among clinical isolates in Wales

We first used short-read sequencing to analyse 540 *Klebsiella* isolates collected between 2007 and 2020 that had largely been referred to SACU at PHW for investigation of antimicrobial resistance or obtained from blood cultures (regardless of the resistance phenotype) (**Supplementary Table 1**). They also included three contextual isolates from UK NEQAS. Overall, 387/540 (71.7%) isolates were from clinical samples (including 238 from blood, 93 from urine, 23 from a wound site, 33 other/unknown), 143/540 (26.5%) from screen samples, 6/540 (1.1%) from environmental samples and 4/540 (0.7%) from an unknown source. The clinical and screen isolates were recovered from 405 patients in healthcare facilities across Wales, of whom 72 (17.8%) contributed two or more isolates (up to eight).

Among our collection, *K. pneumoniae* was the dominant species, accounting for 421/540 (78.0%) isolates. A further eight species were also identified: *K. aerogenes* (33/540; 6.1%), *K. michiganensis* (27/540; 5.0%), *K. variicola* (23/540; 4.3%), *K. oxytoca* (17/540; 3.1%), *K. quasipneumoniae* (11/540; 2.0%), *K. grimontii* (5/540; 0.9%), *K. planticola* (2/540; 0.4%) and *K. pasteurii* (1/540; 0.2%). A phylogenetic tree of the 421 *K. pneumoniae*, constructed from an alignment of 4034 core genes, demonstrated a high number of distinct lineages (**Figure 1;** https://microreact.org/project/aSJqpc9MZVZcurTWtoadW8-k-pneumoniae-phw-n421). We identified 130 STs across this species, with 98 (75.4%) of these restricted to a single patient. However, some STs were observed frequently including ST307 (88 isolates; 55 patients), ST1788 (43 isolates; 30 patients), ST20 (19 isolates; 10 patients), ST15 (19 isolates; 16 patients) and ST14 (17 isolates; 12 patients). Phylogenetic analyses of all other *Klebsiella* species also revealed a high diversity among each, with most isolates occurring as singletons or in small clusters (see **Supplementary Table 3** for Microreact URLs).

**Figure 1.**
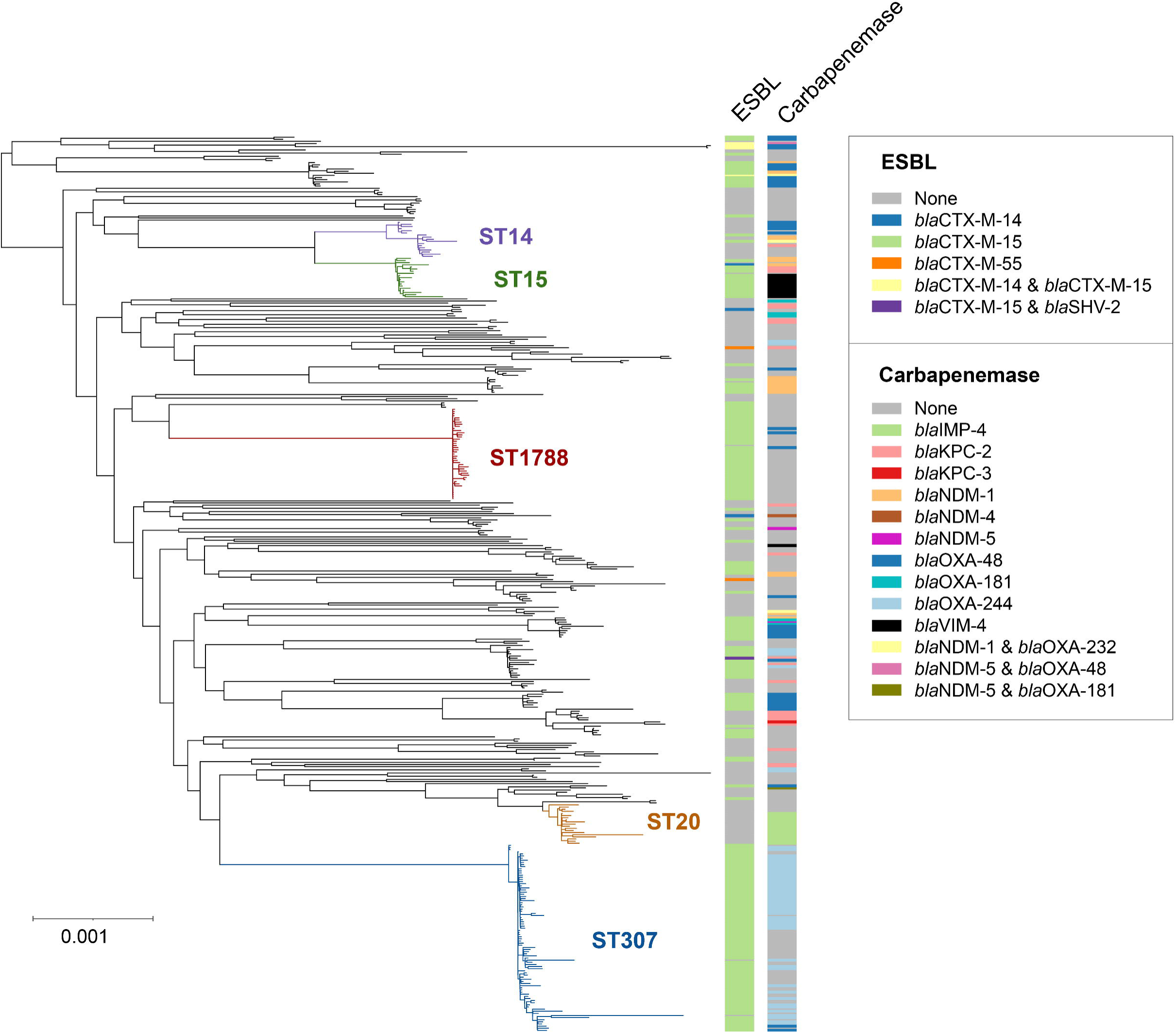
Maximum likelihood phylogenetic tree of 421 *K. pneumoniae* isolates demonstrates high diversity. The tree was constructed from an alignment of 4034 core genes that were present in ≥99% of *K. pneumoniae* isolates referred to the SACU laboratory at PHW. The most frequently observed STs are highlighted. Metadata columns show the ESBL and carbapenemase variants found among the isolates (if applicable). The scale indicates the number of SNPs per variable site. The tree is available to view interactively and with additional metadata at https://microreact.org/project/aSJqpc9MZVZcurTWtoadW8-k-pneumoniae-phw-n421

Inspection of the *Klebsiella* genomes for antimicrobial resistance markers revealed that 282/540 (52.2%) isolates carried an ESBL gene, which was largely *bla*_CTX-M-15_ (228/282; 80.9%) (**Figure 1**). 207/540 (38.3%) isolates carried a carbapenemase gene. The most frequently observed carbapenemases were *bla*_OXA-244_ (58/207; 28.0%), *bla*_OXA-48_ (46/207; 22.2%), *bla*_KPC-2_ (31/207; 15.0%), *bla*_NDM-1_ (29/207; 14.0%), *bla*_IMP-4_ (16/207; 7.7%) and *bla*_VIM-4_ (12/207; 5.8%). Carbapenemase-encoding isolates were found in six of the nine observed species, although *K. pneumoniae* accounted for the vast majority (181/207; 87.4%). Among *K. pneumoniae*, isolates carrying carbapenemase genes were found in 52 STs. However, *bla*_OXA-244_ was mostly carried by ST307 (49/58; 84.5%), *bla*_IMP-4_ by ST20 (15/16; 93.8%), and *bla*_VIM-4_ by ST15 (11/12; 91.7%) or their related variants. Other carbapenemases such as *bla*_KPC-2_ and *bla*_OXA-48_ were more widely dispersed across different species and STs.

### Persistent spread of globally- and locally-important clones of *K. pneumoniae*

Despite the overall high diversity among isolates, our phylogenetic analyses revealed numerous small to large clusters, representing clonal transmission of *Klebsiella* within and between healthcare facilities in Wales. Below we discuss features of the two largest clusters involving *K. pneumoniae* ST307 and ST1788.

### ST307

The most frequently observed ST in our collection, ST307, is an important clone that has been identified in diverse locations worldwide and associated with the ESBL gene, *bla*_CTX-M-15_ (Peirano et al. 2020). In 2019, an outbreak involving ST307 was identified in an acute tertiary hospital in South West Wales (Hospital A), based on epidemiological and resistance profiling data. The isolates showed a multi-drug resistant phenotype (resistance to third-generation cephalosporins, gentamicin, ciprofloxacin +/-ertapenem). The carbapenemase gene, *bla*_OXA-244_, was detected in the subset of isolates that exhibited carbapenem resistance (ertapenem +/-meropenem +/-imipenem). Recognition of the outbreak prompted regular faecal screening for carbapenemase-producing *Enterobacteriaceae* (CPE) carriage in the initial affected ward, and this was extended to additional wards where clinical isolates of *bla*_OXA-244_-producing *K. pneumoniae* were later identified. Here we analysed genome data of 88 isolates from ST307 (or single-locus variants) collected between 2017 and 2020 from ten hospitals and additional GP surgeries in Wales, with particular emphasis on understanding the emergence, evolution and transmission of the outbreak strain.

We constructed a phylogenetic tree of these 88 isolates, also including a further 964 geographically-diverse public ST307 genomes to aid contextualisation. The phylogeny demonstrated that 82/88 (93.2%) of the ST307s from Wales formed a monophyletic cluster (**Figure 2A**; https://microreact.org/project/1ZRxgto9HVsaV6qRHAuf6P-st307s-phw-n1052). This is indicative of a single major introduction of this strain into Wales followed by local spread. The cluster incorporated clinical isolates from 19 patients (of whom 7 had bloodstream infections), as well as screen isolates from a further 31 patients.

**Figure 2.**
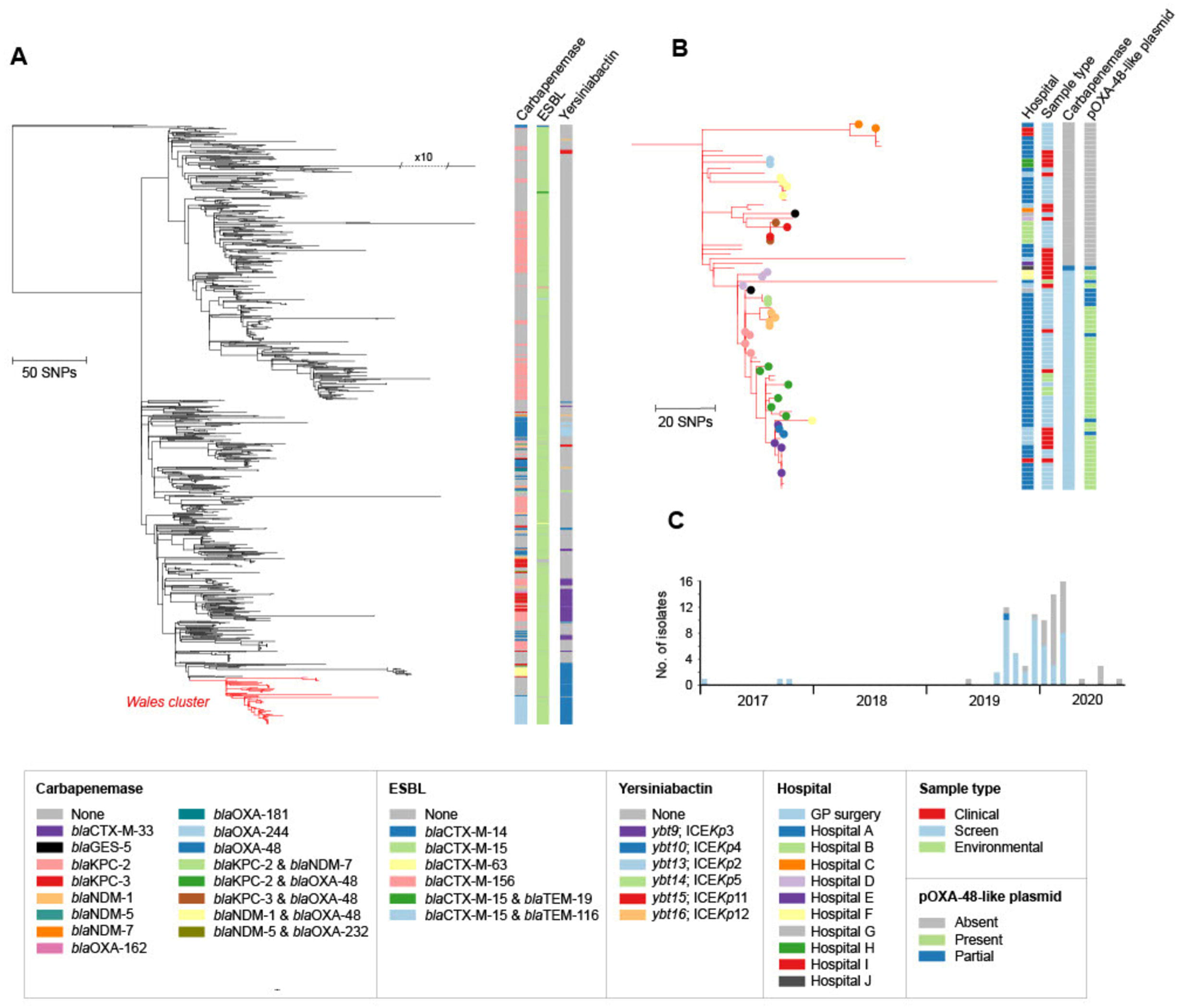
A) Maximum likelihood phylogenetic tree of 1052 *K. pneumoniae* ST307 isolates demonstrates a main introduction of ST307 into Wales. The tree includes 964 ST307 isolates from public archives (from diverse international locations) and 88 isolates from the PHW collection. The tree was constructed from an alignment generated by mapping all short sequence reads to an ST307 reference genome (ARGID_32304) and after the removal of recombined regions. 82 ST307 isolates from the PHW collection form a monophyletic cluster highlighted in red. Metadata columns show the carbapenemase, ESBL and yersiniabactin variants among all isolates (if applicable). The tree is available to view interactively and with additional metadata at https://microreact.org/project/1ZRxgto9HVsaV6qRHAuf6P-st307s-phw-n1052 **B)** Cluster of 82 ST307 isolates from the PHW collection, as zoomed in from (A). Filled circles of the same colour on the tree tips represent isolates from the same patient. Metadata columns show the sampling hospital, sample type, carbapenemase variant and the presence/absence of the pOXA-48-like plasmid. For the latter, the plasmid was scored as “present” if sequence reads mapped to ≥90% of the pOXA48a reference plasmid (accession JN626286), “partial” if between 70% and 90% and “absent” if <70%. The scale bars in (A) and (B) represent the number of SNPs. **C)** Timeline showing the isolation dates of the 82 clustered ST307 isolates from the PHW collection. Bars are coloured by the carbapenemase variant (if applicable).

Almost all of the 82 clustered isolates carried *bla*_CTX-M-15_ (80/82; 97.6%) while a descendent subclade of the cluster later acquired *bla*_OXA-244_ (49/82; 59.8%). A single isolate also carried the combination of *bla*_CTX-M-15_ and *bla*_OXA-48_, the latter of which has a single amino acid substitution relative to *bla*_OXA-244_. We searched for alterations (including truncations) in major outer membrane porin genes that have been shown to reduce susceptibility to carbapenems when acting in concert with ESBL, AmpC or carbapenemase enzymes (Martinez-Martinez et al. 1999; Hamzaoui et al. 2018). We found that all 82 isolates carried a truncated *ompK35* gene while 12/82 (14.6%) also possessed a truncated *ompK36* gene.

Isolates in the *bla*_OXA-244_ subclade were characterised by low pairwise SNP differences (range 0-116; mean 15.5) and mostly collected from patients (both clinical and screen) or environmental samples in Hospital A (40/49; 81.6%) over the outbreak period (**Figures 2B & 2C**). This is consistent with heightened within-hospital transmission. The remaining nine *bla*_OXA-244_-encoding isolates were recovered from other nearby hospitals and GP surgeries in the South Wales region. Notably, the first sampled isolates from this subclade were obtained from a single patient in 2017 (outside of Hospital A). This demonstrates the acquisition of *bla*_OXA-244_ by this strain and emergence of this subclade at least two years prior to its later expansion within Hospital A.

The non-*bla*_OXA-244_-encoding isolates in the Welsh ST307 cluster had been referred to SACU from 2019 onwards, largely prompted by the similarity of their phenotypic profiles to the *bla*_OXA-244_-encoding outbreak isolates rather than by epidemiological links. Despite their collection over a limited timeframe, we found that they were considerably more divergent than the *bla*_OXA-244_-encoding isolates (pairwise SNP differences from 0-134; mean 54.4) (**Figure 2B**). They were also recovered from a wider number of hospitals in South Wales (nine in total) although included isolates from Hospital A. These findings are further consistent with the undetected circulation of this ST307 strain across South Wales over several years prior to the emergence and expansion of the *bla*_OXA-244_-encoding outbreak subclade in Hospital A in 2019.

The *bla*_OXA-244_-encoding ST307 outbreak was declared formally closed in May 2020 with no further isolates of this subclade identified since March 2020.

### ST1788

ST1788 is the second most common ST of *K. pneumoniae* in our collection. In contrast to ST307, this has rarely been observed elsewhere to our knowledge. SACU first detected ST1788 during the 2019 ST307 outbreak investigation and it continued to be recovered across South Wales up until the end of the study period (November 2020). The 43 ST1788 isolates in this sequenced collection are derived from 30 patients (including 18 with clinical isolates and 3 with blood isolates) across seven hospitals and additional GP surgeries. They include 20 isolates from 13 patients obtained from a rehabilitation unit (Hospital B) in South East Wales from early 2020 onwards.

To investigate the evolution and spread of this strain, we constructed a phylogenetic tree of the 43 ST1788 isolates from Wales, together with two publicly available ST1788 genomes from Nigeria. This revealed that the Welsh isolates formed a distinct cluster (**Figure 3;** https://microreact.org/project/rMY2fakRxkNKXqqBPs3is6-st1788-phw-n45). Although additional contextual data are required to draw strong conclusions on the routes of strain spread, these findings are consistent with a single introduction into South Wales followed by local transmission within and between healthcare facilities. This is further supported by the low SNP differences between our isolate pairs (0-36 SNPs, mean 17.9), calculated after recombination removal, suggestive of a recent clonal expansion.

**Figure 3.**
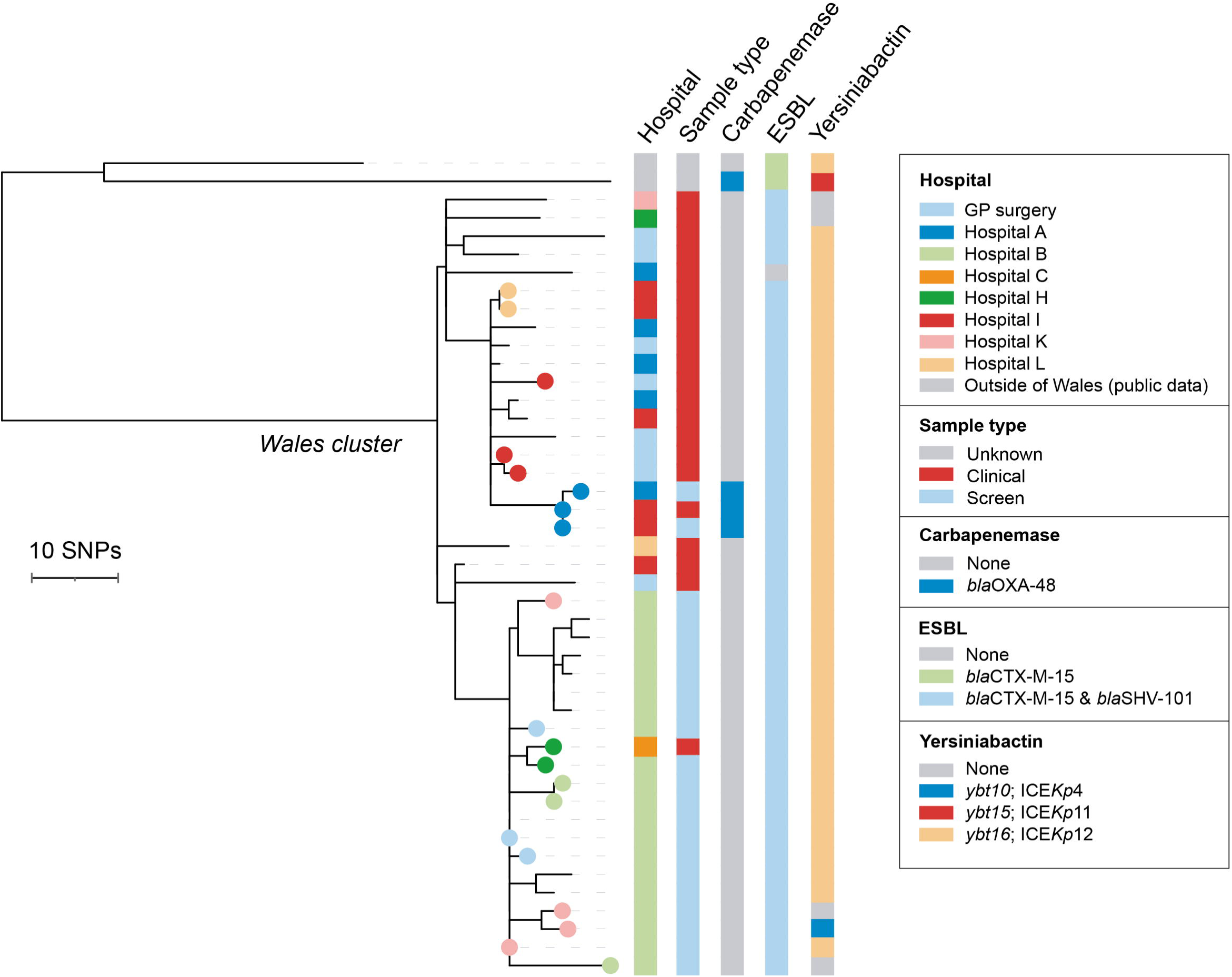
Maximum likelihood phylogenetic tree of 45 ST1788 isolates demonstrates high clonality among isolates from Wales. The tree includes two isolates from public archives (both from Nigeria) and 43 from the PHW collection. It was constructed from an alignment generated by mapping all short sequence reads to an ST1788 reference genome (ARGID_33748) and after the removal of recombined regions. The 43 ST1788 isolates from the PHW collection form a monophyletic cluster. Filled circles of the same colour on the tree tips represent isolates from the same patient. Metadata columns show the sampling hospital, sample type, and carbapenemase, ESBL and yersiniabactin variants (if applicable). The scale bar represents the number of SNPs. The tree is available to view interactively and with additional metadata at https://microreact.org/project/rMY2fakRxkNKXqqBPs3is6-st1788-phw-n45

Our phylogenetic analysis also revealed that the isolates obtained from patients in Hospital B clustered together, also with a bloodstream isolate from an inpatient at an acute tertiary care hospital in South East Wales (Hospital C) who was transferred from Hospital B (**Figure 3**). This demonstrated the occurrence of a single introduction of the strain into Hospital B followed by within-hospital spread. Apart from the isolate obtained from the Hospital C patient, all isolates obtained from Hospital B were screen isolates, obtained via an enhanced screening programme at this site.

Almost all ST1788 isolates from Wales carried *bla*_CTX-M-15_ (42/43; 97.7%) while three isolates from one patient also carried *bla*_OXA-48_ (**Figure 3**). The majority (39/43; 90.7%) had intact *ompK35* and *ompK36* genes. Notably, all ST1788 isolates possessed the KL2 capsule type, which has been associated with increased virulence (Shon et al. 2013), and most (39/43; 90.7%) also possessed a yersiniabactin locus.

At the time of writing, transmission of ST1788 remains an ongoing concern, in particular in Hospital B. We also have evidence of ongoing secondary cases at nearby hospitals within the health board, linked by epidemiology and genomic typing, and have recently recovered ST1788 isolates from patients in other regions of Wales.

### Plasmid epidemiology of *bla*_KPC-2_-producing *Klebsiella* reflects distinct hospital networks and referral patterns between North and South Wales

We next aimed to elucidate the spread of particular carbapenemase genes in Wales where we could not identify clonal transmission as a key driver. We first investigated *bla*_KPC-2_ genes, which were found in 30 isolates from five different species among our collection (excluding a NEQAS isolate with *bla*_KPC-2_). While the majority were from *K. pneumoniae* (21/30; 70.0%), 13 different STs from this species were represented and no single ST accounted for more than four isolates.

Using our short-read data with the TETyper tool (Sheppard et al. 2018), we first determined whether the *bla*_KPC-2_ genes were carried on the Tn*4401* transposon, which has previously been associated with *bla*_KPC_ gene spread (Naas et al. 2008; Cuzon et al. 2011). In ten isolates (10/30; 33.3%), TETyper did not identify any known Tn*4401* variant. However, the remaining isolates (20/30; 66.7%), from a variety of species and STs, carried *bla*_KPC-2_ on the Tn*4401a* variant. These were recovered from 16 patients between 2013 and 2020. Tn*4401a* is the same variant associated with endemic spread of *bla*_KPC-2_-encoding *Enterobacteriaceae* in North-West England recognised from 2008 onwards (Stoesser et al. 2020). Notably, the majority (17/20; 85.0%) of our isolates with *bla*_KPC-2_ carried on Tn*4401a* were sampled from patients in North Wales. These findings likely reflect acquisition of *bla*_KPC-2_-encoding strains and/or plasmids in North-West England hospitals, consistent with the typical referral of patients in North Wales to this region for specialist treatment.

To further elucidate the diversity of plasmids carrying *bla*_KPC-2_ on Tn*4401a*, we long-read sequenced 13 of these isolates (including the NEQAS isolate) (**Supplementary Table 2**). These were selected to represent the diversity of species and STs observed. We obtained complete plasmids carrying *bla*_KPC-2_ from hybrid (combined short- and long-read) assemblies in all 13 isolates. One *K. pneumoniae* ST11 isolate (ARGID_32407) also carried an additional copy of *bla*_KPC-2_ on the chromosome. Of the 13 *bla*_KPC-2_ plasmids, all possessed either one or two IncF replicons, sometimes in combination with other replicon types (e.g. IncR). They ranged in size from 113.5-210.3kb. In particular, we found that seven had a similar structure and size to pKpQIL-like plasmids, a family of plasmids represented by pKpQIL (Leavitt et al. 2010).

A visual comparison of pKpQIL-like plasmids from our collection, together with two published pKpQIL-like plasmids from North-West England (Stoesser et al. 2020) and pKpQIL (Leavitt et al. 2010), showed the high structural conservation overall (**Figure 4**). However, two plasmids (from ARGID_31921 and ARGID_24826) had lower nucleotide similarity with pKpQIL over one half, likely as a result of recombination. All others showed high nucleotide similarity to pKpQIL, differing by 11-32 SNPs. We also found some support for recent common ancestry between plasmids from North Wales and North-West England. For example, the plasmids from ARGID_18508 and ARGID_31165 (both North Wales) differed from trace384 (North-West England) by only 4 SNPs.

**Figure 4.**
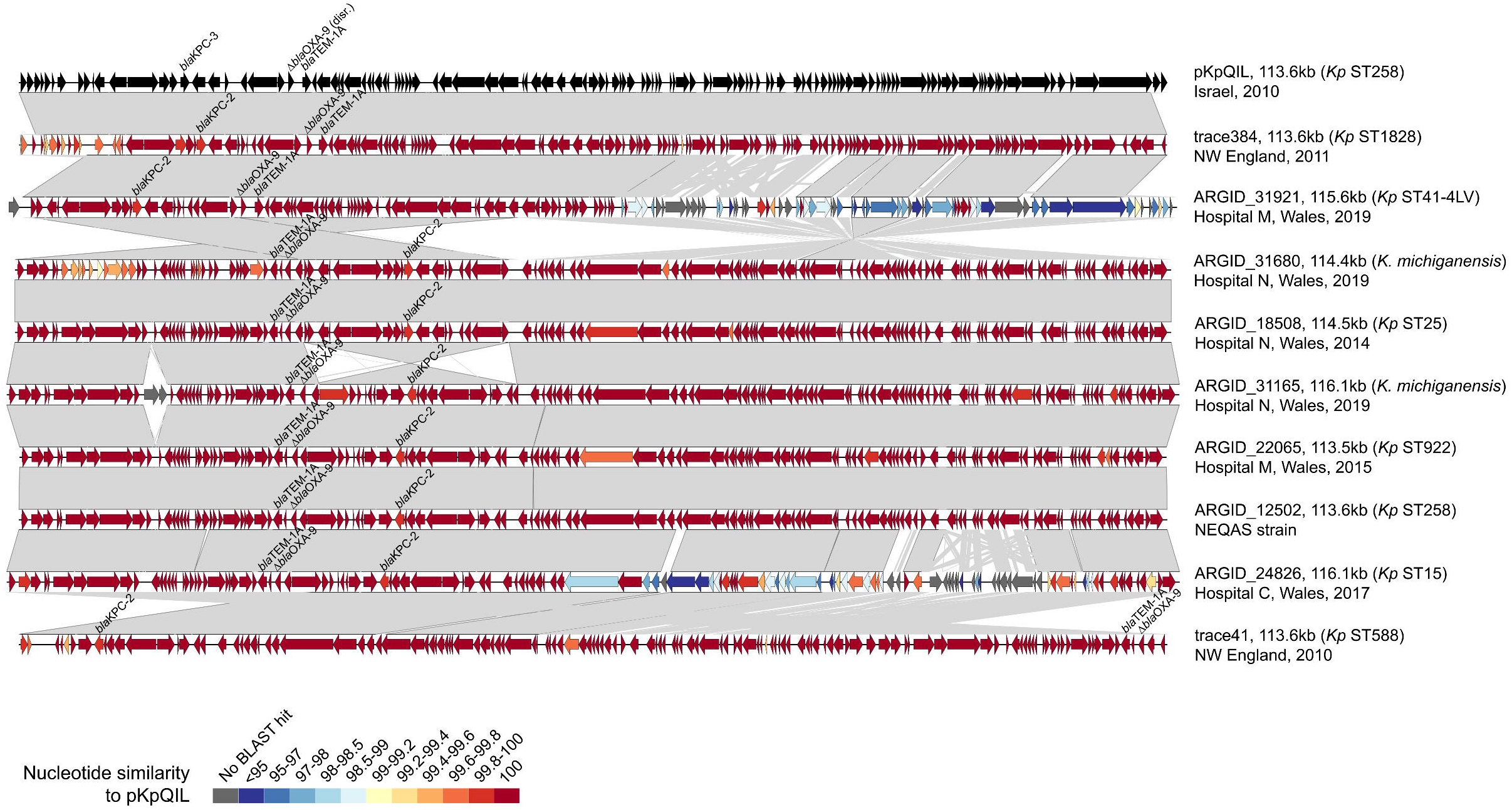
Pairwise comparisons between complete *bla*_KPC_-encoding pKpQIL-like plasmids demonstrate high structural and sequence conservation. The plasmids include the original pKpQIL sequence (accession GU595196) (Leavitt et al. 2010), seven from the PHW collection and two previously sequenced plasmids recovered in North-West England hospitals (Stoesser et al. 2020). Open reading frames (ORFs) are depicted as arrows with resistance genes (including *bla*_KPC_) labelled. Grey blocks between plasmid pairs indicate regions of homology. The colour of the ORFs represents the percentage nucleotide similarity to pKpQIL via BLASTn.

We next assessed the remaining ten *bla*_KPC-2_-carrying isolates with no known Tn*4401* variant. These isolates were from five patients and one environmental sample, and from a mixture of species and STs. They were recovered between 2017 and 2020 from three hospitals in South Wales, with seven isolates from Hospital A. Inspection of the short-read genome assemblies demonstrated a shared genetic context around *bla*_KPC-2_. We also found that all ten isolates had an IncN replicon in common. To investigate the possibility of *bla*_KPC-2_-IncN plasmid spread, we long-read sequenced six of these isolates from different species and/or STs (**Supplementary Table 2**).

We obtained a complete IncN plasmid from the hybrid assemblies of all six isolates, and confirmed carriage of *bla*_KPC-2_ in a conserved genetic environment (next to *bla*_TEM-1_) on these plasmids. The IncN plasmids varied considerably in size from 48.2kb to 137.4kb, and the largest also carried an additional IncFII replicon. Visual comparison of the plasmids showed that they shared high nucleotide similarity across shared “core” regions (**Figure 5**). This is despite frequent recombination events leading to alternate structural arrangements and a large variation in “accessory” sequence. We also found high nucleotide similarity among the shared regions of these plasmids and a previously described IncN plasmid from an ST307 isolate recovered in the UK (accession KY271414) (Villa et al. 2017) (**Figure 5**). These findings point to the recent emergence and spread of this *bla*_KPC-2_-encoding IncN backbone between different *Klebsiella* species and STs in South Wales and the potentially wider UK region.

**Figure 5.**
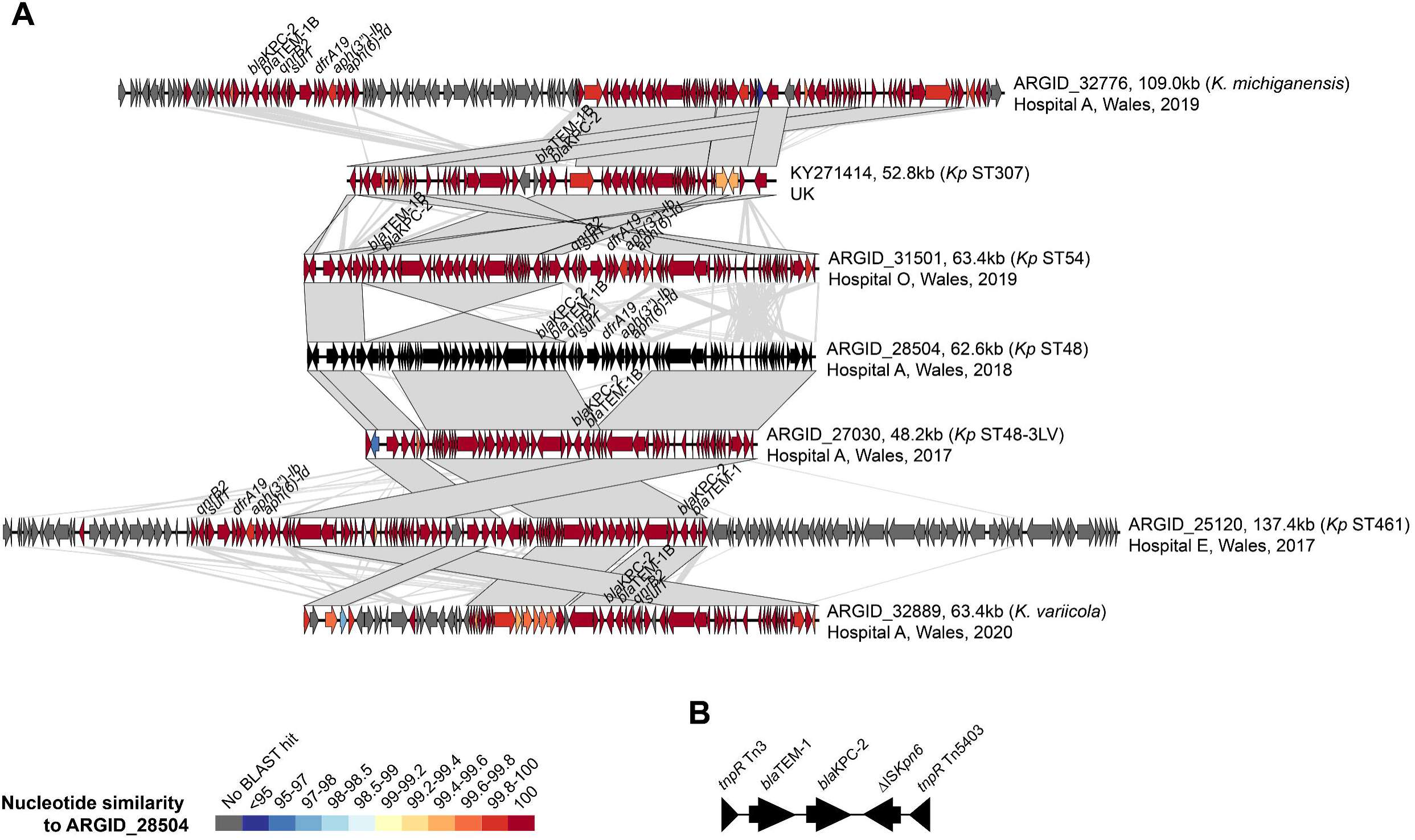
A) Pairwise comparisons between complete *bla*_KPC-2_-encoding IncN plasmids demonstrate substantial plasticity despite high nucleotide similarity among shared backbone regions. The plasmids include a previously published sequence (accession KY271414) (Villa et al. 2017) and six from the PHW collection. ORFs are depicted as arrows with resistance genes (including *bla*_KPC-2_) labelled. Grey blocks between plasmid pairs indicate regions of homology. The colour of the ORFs represents the percentage nucleotide similarity to the plasmid from ARGID_28504 via BLASTn. **B)** Conserved genetic environment of *bla*_KPC-2_ among the IncN plasmids.

### Frequent carriage and horizontal dissemination of *bla*_OXA-48-like_ genes via pOXA-48-like plasmids and related variants

The most common type of carbapenemase genes among our *Klebsiella* collection were *bla*_OXA-48-like_ genes, which comprised *bla*_OXA-244_ (*n*=58), *bla*_OXA-48_ (*n*=46), *bla*_OXA-181_ (*n*=7) and *bla*_OXA-232_ (*n*=3) variants. While most *bla*_OXA-244_-encoding isolates (49/58; 84.5%) belonged to the ST307 lineage, other *bla*_OXA-48-like_ genes were identified in diverse species and STs. In particular, *bla*_OXA-48_ genes were found in three species (*K. pneumoniae, K. quasipneumoniae, K. oxytoca*) and 15 different STs of *K. pneumoniae*, and recovered between 2013 and 2020.

We investigated a possible association of *bla*_OXA-48-like_ genes with pOXA-48-like plasmids, which are highly conserved ∼61-63kb IncL plasmids shown previously to frequently carry these carbapenemase genes (Poirel et al. 2012; David et al. 2020). To do this, we mapped the short reads of all isolates in our collection (including those without *bla*_OXA-48-like_ genes) to the 61.8kb pOXA48a reference plasmid (Poirel et al. 2012). 113/540 (20.9%) isolates mapped to ≥70% of the plasmid length (including 95 that mapped to ≥90%), suggesting that they harboured the same or a similar plasmid. Most of the remaining isolates mapped to ≤10%. The 113 isolates with ≥70% plasmid mapping were from 61 different patients in 15 hospitals and additional GP surgeries, largely from across South Wales. They accounted for most isolates with *bla*_OXA-48-like_ genes, including all of those with *bla*_OXA-244_ (58/58) and *bla*_OXA-48_ (46/46), and 1/3 (33.3%) of those with *bla*_OXA-232_. Only 8/113 (7.1%) possessed no *bla*_OXA-48-like_ gene. Among the 113 isolates, 94 (83.2%) possessed an IncL replicon (the same as pOXA48a). We did not find an IncL replicon in the remaining 19/113 (16.8%) isolates, but instead found either an IncM1 *(n*=18) or IncM2 (*n*=1) replicon. IncL and IncM plasmids are known to be closely related and indeed were previously classified in the same family, IncL/M (Carattoli et al. 2015). However, the latter have not been implicated as vectors of *bla*_OXA-48-like_ genes to date.

Analysis of SNP diversity among the 113 mapped plasmid sequences demonstrated 0-5 differences among many (≥75%) of the sequences, despite their recovery from diverse locations and times. This high similarity was reflected in a minimum spanning tree (**Figure 6A**). Only some sequences were considerably more divergent, including those associated with IncM replicons (**Figure 6B**). While the overall low amount of SNP diversity among these plasmid sequences makes inference of transmission events difficult, we did find identical plasmid sequences shared among different species and STs, suggestive of recent horizontal spread. For example, we found 0 SNPs among 55 sequences (all associated with *bla*_OXA-244_), which included 48 from the *K. pneumoniae* ST307 outbreak isolates, two from ST1430, two from ST48, two from ST353 and one from *K. oxytoca* (**Figure 6C**). The vast majority of these, including the non-ST307s, were recovered from Hospital A, supporting the possibility of horizontal transmission. Notably, five of these non-ST307s were recovered in 2017-2018, implying local circulation of this plasmid variant prior to the expansion of *bla*_OXA-244_-encoding ST307 in 2019.

**Figure 6.**
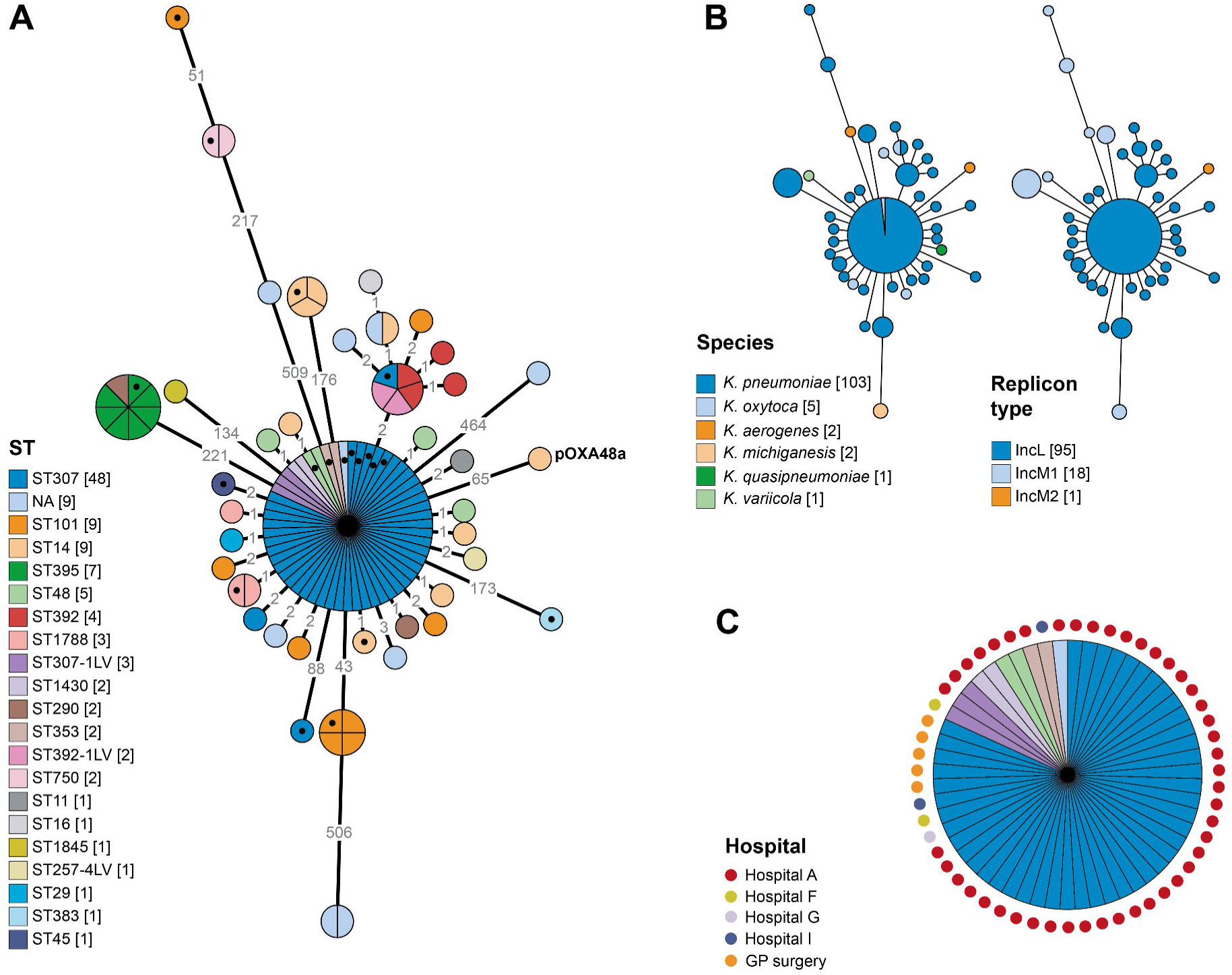
A) Minimum spanning tree of 113 pOXA-48-like plasmid sequences demonstrates high nucleotide conservation as well as some divergent sequences. Plasmid sequences included in the tree, which were obtained by mapping short reads to pOXA48a (accession JN626286) (Poirel et al. 2012), are from isolates with ≥70% mapping to this reference sequence. The tree was constructed from a SNP distance matrix. The branch lengths are displayed using a log scale and indicated on the branches. The nodes are partitioned by (and their sizes proportional to) the number of represented isolates. Isolates are coloured by ST with the number of isolates from each ST indicated in square brackets. Isolates marked with a black filled circle were selected for long-read sequencing to enable complete plasmid sequencing. **B)** The same trees as in (A) coloured by species of the host isolate or replicon type identified from the genome. **C)** The major node from the tree in (A) coloured by ST, with annotations marking the sampling hospitals.

Finally, we used the short-read data to investigate structural differences among the 113 mapped plasmid sequences. To do this, we calculated and visualised the short-read coverage across the length of pOXA48a, relative to the median chromosomal coverage of the host strain (**Figure 7**). This revealed that nine of the *bla*_OXA-244_-encoding ST307 outbreak isolates lacked coverage across parts of the plasmid (8.6-16.6kb) that include the *tra* region in most cases, with a conserved breakpoint at one end of the deletion. This suggests that these plasmids may have lost the ability to self-conjugate and/or be mobilised. Notably, they did not cluster into a single clade in the core genome-based phylogenetic tree, suggesting that the deletions occurred on multiple independent occasions (**Figure 2**).

**Figure 7.**
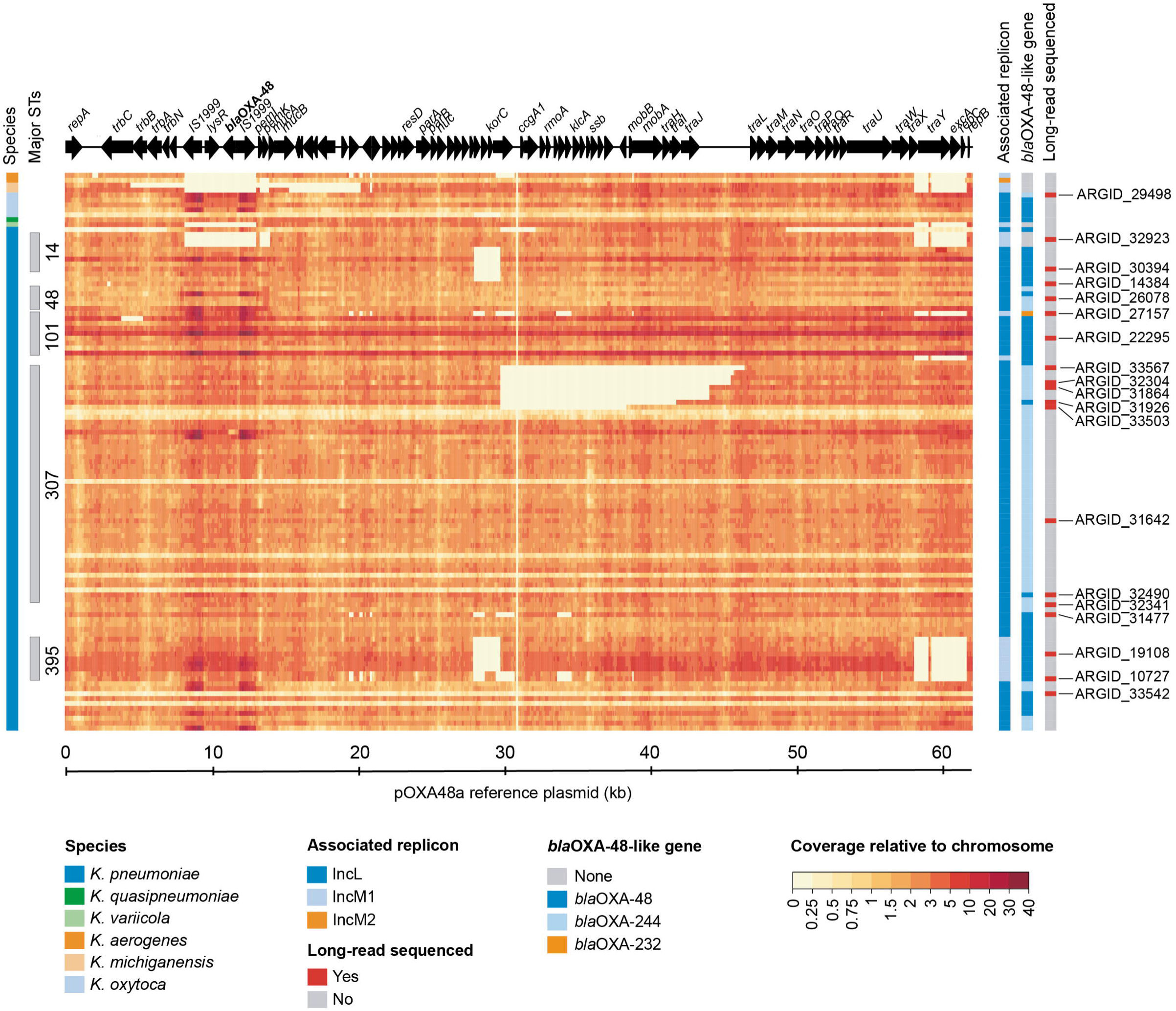
Heatmap showing the short-read coverage across the pOXA48a reference sequence indicates deletions among some isolates, including several involving the *tra* region in ST307. The plasmid coverage is shown relative to the median chromosomal coverage for 113 isolates with ≥70% mapping to pOXA48a (accession JN626286) (Poirel et al. 2012). ORFs from the pOXA48a plasmid are depicted at the top, including *bla*_OXA-48_. Metadata columns indicate the host species, those belonging to frequently observed STs, and replicon and *bla*_OXA-48-like_ variants identified from the isolate genomes. The far-right column indicates isolates selected for long-read sequencing to enable complete plasmid assembly.

### Structural evolution of pOXA-48-like plasmids including repeated loss of conjugative ability in *K. pneumoniae* ST307

We next used our short-read analysis of the SNP and structural diversity among pOXA48-like plasmids to guide selection of 19 isolates for further investigation with long-read sequencing (**Supplementary Table 2**). These were chosen to investigate: a) the possible carriage of *bla*_OXA-48-like_ genes on IncM plasmids and their structural differences with IncL plasmids; b) other divergent variants of pOXA-48-like plasmids and c) the evolution of pOXA-48-like plasmids in the ST307 outbreak lineage. We constructed hybrid assemblies and obtained complete IncL or IncM1 plasmids in all 19 isolates, which ranged in size from 46.8kb to 73.0kb.

Among the 19 complete plasmids, we obtained three with an IncM1 replicon and confirmed carriage of *bla*_OXA-48_ genes on these. Overall they showed high structural homology to the IncL plasmids albeit with key differences (**Figure 8**). These included a section with the replicon typing determinants that lacks homology, as well as lower sequence similarity over the *tra* gene cluster region. However, large parts of the IncL and IncM1 plasmids shared 100% nucleotide similarity, including the region containing *bla*_OXA-48_, suggestive of recent recombination.

**Figure 8.**
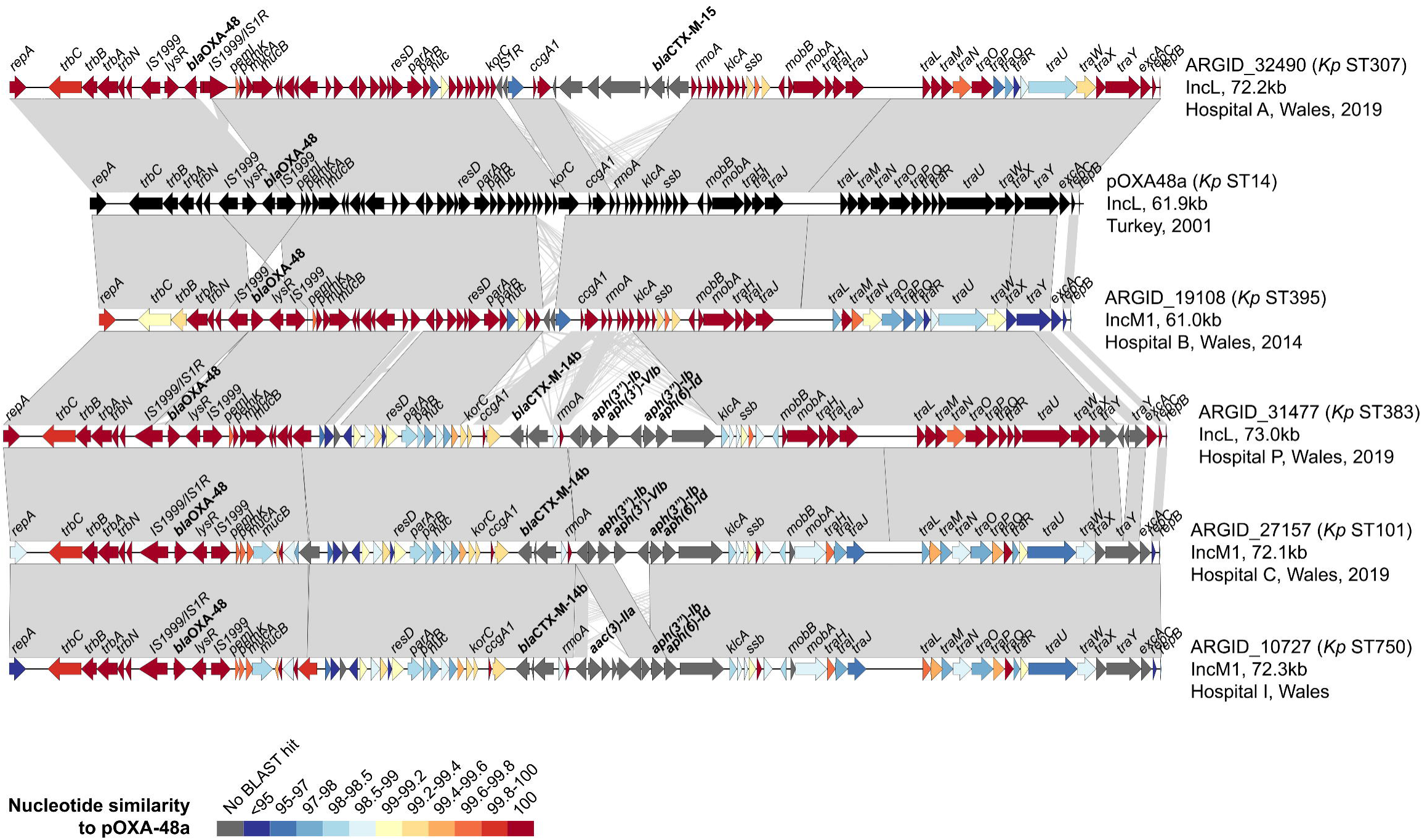
Pairwise comparisons between complete pOXA-48-like plasmids show structural differences to the “classic” backbone of pOXA48a, including insertion of islands carrying resistance genes. The plasmids include pOXA48a (accession JN626286) (Poirel et al. 2012) and five from the PHW collection. They are from a range of STs and include those with both IncL and IncM1 replicon types. ORFs are depicted as arrows with resistance genes (including *bla*_OXA-48_) labelled. Grey blocks between plasmid pairs indicate regions of homology. The colour of the ORFs represents the percentage nucleotide similarity to pOXA48a via BLASTn.

Among other isolates selected for long-read sequencing based on the high structural and/or SNP divergence of their pOXA-48-like plasmids from pOXA48a, we found several with novel integrations (**Figure 8**). These included an island harbouring *bla*_CTX-M-15_ in one plasmid recovered from a *K. pneumoniae* ST307 strain (unrelated to the ST307 outbreak strain). Furthermore, a genetic island with another ESBL gene, *bla*_CTX-M-14b_, as well as aminoglycoside resistance genes, was found in both IncL and IncM1 plasmids recovered from *K. pneumoniae* ST101, ST383 and ST750 isolates.

Finally, we analysed complete pOXA-48-like plasmids from six ST307 outbreak-associated isolates to investigate the putative deletions involving the *tra* region. These included ARGID_31642, in which no deletion had been observed, and five isolates with putative deletions. As expected, we found that the plasmid from ARGID_31642 had a similar structure and size to pOXA48a (**Figure 9**). All others possessed a deletion with a conserved breakpoint at one end adjacent to an IS*1R* element. The variable size and nature of these deletions further confirmed that they had occurred on multiple independent occasions. This suggests that the deletions confer a fitness advantage. The deletions involved one or more *tra* genes and the *oriT* region in four of the five cases, with one plasmid (from ARGID_33503) retaining these regions despite a deletion. We used conjugation experiments to test the ability of the six isolates with complete pOXA-48-like sequences to transfer the plasmid to both *E. coli* (J53 strain) and *K. pneumoniae* (71.1 strain), using two different ratios of donor/recipient (1:3 and 5:1). These confirmed that both ARGID_31642 and ARGID_33503 could transfer the plasmid, while no transfer of plasmids with deletions in the *tra*/*oriT* regions was observed (**Supplementary Table 4**).

**Figure 9.**
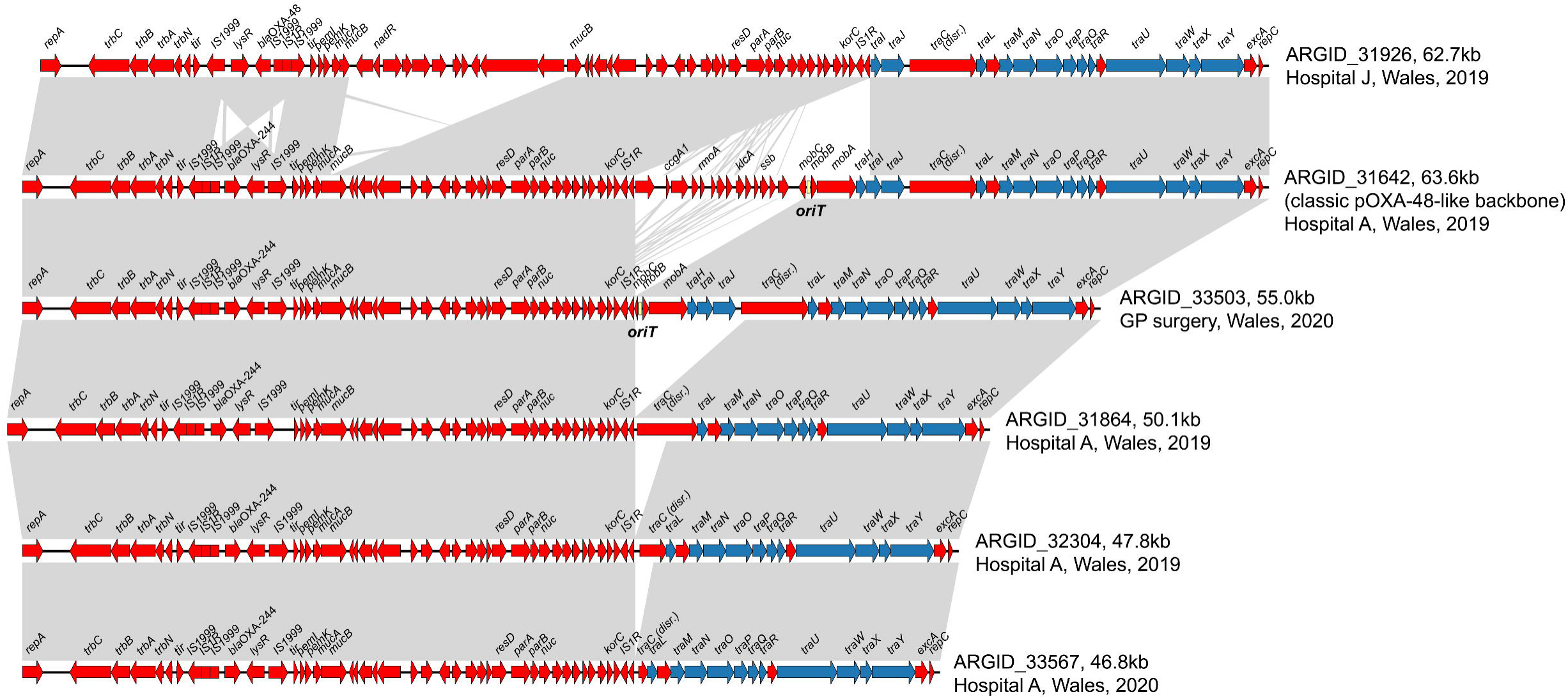
Pairwise comparisons between complete pOXA-48-like plasmids from ST307 demonstrate multiple independent deletions of regions adjacent to an IS*1R* element. One plasmid (from ARGID_31642), which possesses the “classic” pOXA-48-like backbone and which lacks these deletions, is included. ORFs are depicted as arrows with resistance genes (including *bla*_OXA-48_/*bla*_OXA-244_) labelled. Grey blocks between plasmid pairs indicate regions of homology. ORFs coloured blue are predicted to be involved in conjugation while those in red have other predicted functions. The *oriT* region is indicated if identified in the plasmid.

## Discussion

In line with other geographic regions worldwide, *Klebsiella* infections represent a substantial healthcare burden in Wales and are the third most common cause of bloodstream infections (PHW, 2018). While rates of resistance to carbapenems remain low (<1% among bloodstream isolates in 2017) (PHW, 2018), the rapid global rise of resistance necessitates vigilant monitoring (Cassini et al. 2019; Castanheira et al. 2019). Here we describe the major strains and plasmids driving spread of multi-drug resistant *Klebsiella* in Wales between 2007 and 2020, providing a detailed view of resistance dynamics in a largely non-endemic setting.

We found that more than half of carbapenemase-producing *Klebsiella* isolates (124/207; 59.9%) referred to our laboratory at PHW belonged to just nine STs from *K. pneumoniae*. This finding echoes numerous other studies reporting a concentration of resistant *Klebsiella* in “high-risk” STs (David et al. 2019; Bonnin et al. 2020; Di Pilato et al. 2021). The major carbapenemase-producing clones detected in our region include the globally important *K. pneumoniae* strains, ST307, ST20 and ST15, which carried *bla*_OXA-244_, *bla*_IMP-4_ and *bla*_VIM-4_, respectively. We also identified spread of a strain which has rarely been reported elsewhere, ST1788, carrying *bla*_CTX-M-15_ and which also acquired *bla*_OXA-48_ in one patient. Detailed phylogenetic analyses of the ST307 and ST1788 clones revealed that a single major introduction of each into Wales, followed by local spread within and between hospitals, accounted for the dominance of these clones among patient isolates.

Our phylogenetic analyses also suggested that the ST307 strain had been circulating in Wales for several years prior to its acquisition of *bla*_OXA-244_ and an outbreak involving the *bla*_OXA-244_-encoding subclade centred on Hospital A in 2019. The lack of recognition of ST307 by SACU until the 2019 outbreak reflects the lack of active surveillance or typing of third-generation cephalosporin-resistant *Klebsiella* infections in Wales over this time period, and the absence of clear epidemiological links between infections. Since the outbreak involving the *bla*_OXA-244_-encoding subclade was declared closed, it is possible that the ST307 strain (without *bla*_OXA-244_) continues to disseminate and be responsible for a significant proportion of invasive infections. However, in the absence of epidemiological links and active surveillance of all isolates, this once more remains undetected. Yet past reports demonstrate the ability of ST307 to readily acquire additional resistance determinants and expand rapidly (Lowe et al. 2019; Haller et al. 2019). Thus, as our laboratory and others move towards using WGS data for routine surveillance of *Klebsiella*, this experience demonstrates the importance of urgently tailoring our systems to detect and control the spread of high-risk clones, ideally prior to acquisition of additional resistance determinants.

Our analysis also revealed the dominant plasmid vectors driving spread of *bla*_KPC-2_ and *bla*_OXA-48-like_ genes in Wales, which were found among diverse species and strain backgrounds. We found that *bla*_KPC-2_ genes from North and South Wales were largely encoded on different plasmid backgrounds, consistent with the referral of patients from these regions to different hospitals for their tertiary medical care (i.e. to North-West England and South Wales, respectively). Notably, *bla*_KPC-2_ genes from isolates in North Wales collected from 2013 onwards were typically encoded within the Tn*4401a* transposon on IncF plasmids, the same genetic background identified among *bla*_KPC-2_ isolates from North-West England. This region has experienced endemic spread of *bla*_KPC-2_-encoding *Enterobacteriaceae* since 2008, driven largely by horizontal exchange of plasmids and plasmid fragments between strains and species (Stoesser et al. 2020). Thus our findings likely reflect the outward expansion of this complex, multi-species, plasmid-driven outbreak to Wales via hospital networks.

By contrast, *bla*_KPC-2_ genes found among isolates from South Wales since 2017 were typically harboured by IncN plasmids. *bla*_KPC-2_-encoding IncN plasmids have been only rarely reported (Villa et al. 2017) and likely represent an emerging plasmid background transmitting these genes. Despite their large variation in size and structure, the high nucleotide similarity found among their shared regions strongly suggests the existence of a recent common ancestor and thereby the occurrence of local plasmid transmission events between different species and STs in Welsh hospitals. This genomic evidence is further supported by the spatial clustering of these plasmids (i.e. 7/10 were recovered from a single hospital). As exemplified with these IncN plasmids, the large structural variability among some plasmid types (despite recent common ancestry) heightens the challenge of routinely identifying plasmid transmissions, even with complete plasmid sequences. We propose that calculating nucleotide similarity across individual segments (or sliding windows) of plasmid sequences, rather than determining an average over the entire sequence, is crucial for detecting transmissions involving plasmids that can rapidly acquire or lose accessory regions, and/or exchange modular regions with other backbone types.

We found that most *bla*_OXA-48-like_ genes (105/114; 92.1%) encoded by isolates referred to our laboratory were harboured by pOXA-48-like plasmids. These accounted for the clonal spread of *bla*_OXA-244_ by the ST307 outbreak strain and also contributed to the horizontal spread of both *bla*_OXA-48_ and *bla*_OXA-244_ variants across numerous species and STs. Other countries and regions have reported a similar dominance of these plasmids, which have been found to be remarkably conserved over time and space (Skalova et al. 2017; David et al. 2020). Given their importance, we propose that the identification and typing of pOXA-48-like plasmids by genomic surveillance systems would have large gains for detection and control of resistance spread. Due to the high structural conservation of this plasmid family, this is largely achievable using short-read approaches (e.g. mapping or plasmid MLST (Brehony et al. 2019)).

Our analysis did, however, reveal a small proportion of novel divergent variants of pOXA-48-like plasmids. These included those carrying additional resistance genes as well as those associated with IncM, rather than IncL, replicons. We also detected deletions among pOXA-48-like plasmids in the ST307 outbreak lineage which extended to the *oriT* and *tra* regions in most cases. Experiments confirmed that the plasmids lacking these regions could no longer be transferred via conjugation to a recipient *K. pneumoniae* or *E. coli* strain. Since these deletions occurred multiple times independently, we propose that they represent adaptation of the pOXA-48-like plasmid to carriage by ST307 strains, likely via a reduction in the fitness cost imposed on the host.

Altogether this study provides a comprehensive picture of resistance gene spread among *Klebsiella* species in Wales, mediated via both clonal and horizontal transmission. It provides important contextual data for ongoing surveillance as well as key considerations for the design and optimisation of routine genomic surveillance systems in public health laboratories.

## Supporting information

Supplemental material summary and Table 3

## Authors and contributors

SD, MM, KS, MW and LJ conceived the study. SD performed the bioinformatic data analysis and drafted the manuscript. MM, KS, EP, LG and JW performed the sequencing, laboratory experiments and data analysis. OBS, DMA, MW and LJ supervised the study. All authors reviewed and edited the manuscript.

## Acknowledgements

We thank the Pathogen Informatics team at the Wellcome Sanger Institute for informatics support; the Pathogen Genomics Unit laboratory and bioinformatics teams at Public Health Wales; the team at the Specialist Antimicrobial Chemotherapy Unit at Public Health Wales; the Infection Prevention & Control departments for Health Boards, especially those teams which commented on the local outbreaks reported herein; and referring Microbiology Laboratories from across Wales and their host Health Boards.

## Supplementary Tables

**Supplementary Table 1**. Metadata, assembly statistics and genotyping results for 540 short-read sequenced *Klebsiella* isolates included in this study.

**Supplementary Table 2**. Metadata and hybrid assembly statistics for 38 long-read sequenced *Klebsiella* isolates included in this study.

**Supplementary Table 3**. URLs for interactive Microreact projects containing phylogenetic analyses for each *Klebsiella* species (those with ≥5 isolates only) together with all metadata and genotypic data.

**Supplementary Table 4**. Conjugation frequencies determined from mating cultures comprising ST307 donor isolates harbouring a *bla*_OXA-48-like_-encoding pOXA-48-like plasmid and a recipient *K. pneumoniae* or *E. coli* strain.

