## Supplemental material summary and Table 3 for "Genomic surveillance of carbapenem-resistant *Klebsiella* in Wales reveals persistent spread of *K. pneumoniae* ST307 and adaptive evolution of pOXA-48-like plasmids"

**Supplementary Table 1. Provided in Excel spreadsheet.**

**Supplementary Table 2. Provided in Excel spreadsheet.**

**Supplementary Table 3.** URLs for interactive Microreact projects containing phylogenetic analyses for each *Klebsiella* species (those with ≥5 isolates only) together with all metadata and genotypic data.

| **Species** | **Microreact URL** |
| --- | --- |
| *K. pneumoniae* | https://microreact.org/project/aSJqpc9MZVZcurTWtoadW8-k-pneumoniae-phw-n421 |
| *K. aerogenes* | https://microreact.org/project/dsACmepLnijN2GKy9dr29s-k-aerogenes-phw-n33 |
| *K. michiganensis* | https://microreact.org/project/8UDw4RarAEnVkMu4jDzCe5-k-michiganensis-phw-n27 |
| *K. variicola* | https://microreact.org/project/4Ctdt1YhyrPcei6cswD4iw-k-variicola-phw-n23 |
| *K. oxytoca* | https://microreact.org/project/f5w1sF46NFkExbHTLgDiNm-k-oxytoca-phw-n17 |
| *K. quasipneumoniae* | https://microreact.org/project/aER2kfzsHKbin1ZWBvwU3y-k-quasipneumoniae-phw-n11 |
| *K. grimontii* | https://microreact.org/project/jfHhJfsHs244sxb6ZqNziA-k-grimontii-phw-n5 |

**Supplementary Table 4. Provided in Excel spreadsheet.**
